# Allosteric inhibitors of Zika virus NS2B-NS3 protease targeting protease in super-open conformation

**DOI:** 10.1101/2022.03.09.483711

**Authors:** Ittipat Meewan, Sergey A. Shiryaev, Chun-Teng Huang, Yi-Wen Lin, Chiao-Han Chuang, Alexey V. Terskikh, Ruben Abagyan

## Abstract

Zika virus (ZIKV) is a member of the Flaviviridae family and is considered a major health threat causing cases of microcephaly in newborns and Guillain-Barré syndrome in adults. Here we targeted a transient deep and hydrophobic pocket of the super-open conformation of ZIKV NS2B-NS3 protease to overcome the limitations of the orthosteric inhibitors. After virtual docking screening of approximately 7 million compounds against the novel allosteric site we selected the top seven candidates and tested them in an enzymatic assay. Six out of seven top candidates selected by the docking screen inhibited ZIKV NS2B-NS3 protease proteolytic activity at low micromolar concentrations, as well as suppressing viral replication. These six compounds, targeting the selected protease pocket conserved in ZIKV as well as several other Flaviviruses, have opened an opportunity for a new kind of drug candidate that might be useful to treat several flaviviral infections.

## INTRODUCTION

The Zika virus (ZIKV) is an emerging global pathogen declared as a Public Health Emergency of International Concern by the World Health Organization. Approximately 84,000 cumulative cases have been reported globally since 2016 (1–4). The virus is transmitted by the mosquito species *Aedes aegypti* and *Aedes albopictus*, as well as directly from human to human (5–8). The main symptoms of a ZIKV infection including fever, rash, and conjunctivitis are relatively mild and frequently asymptomatic in adults. However, ZIKV may cause microcephaly in children born by mothers infected with ZIKV during the pregnancy (9–11). There is also evidence that ZIKV infection may be linked with Guillain-Barré syndrome, associated with autoimmune degeneration of the myelin sheath in adult’s peripheral neurons (12–14). *Currently, there are no effective drugs against ZIKV or any other flaviviral infection* (15). Furthermore, vaccines against several Flaviviral infections may be problematic due to antibody dependent enhancement phenomenon, with the notable exceptions of Japanese Encephalitis and Yellow Fever (16–17).

The two-part ZIKV protease is essential for viral replication, hence it was considered a potential target for treatment. ZIKV non-structural NS3 protein is a multi-functional protein with protease and helicase activities. The C-terminal 440 residue region of NS3 encodes NS3 helicase (NS3hel) that is involved in RNA replication and RNA capping (18–22). A central cytosolic ~50 residue fragment of the transmembrane NS2B protein, forms a functional NS2B-NS3pro complex with N-terminal ~170-residues of NS3, responsible for the cleavage of ZIKV polyprotein (Figure 1). The cleavage results in activating capsid (C), pre-membrane (pr), membrane (M), envelope (E) structural, and nonstructural (NS1, NS2A, NS2B, NS3, NS4A, NS4B, and NS5) proteins (23–24). NS2B cofactor is required for the protease activity of NS3pro. ZIKV NS2B-NS3pro possesses high proteolytic activity but becomes enzymatically inactive if NS2B cofactor is deleted (25–26). In addition to the viral polyprotein processing, the presence of multiple copies of active viral protease inside of the host cell cytoplasm may lead to the irreversible damage of numerous host cell proteins. Thus, NS2B-NS3pro is a promising drug target for the treatment of ZIKV infection (15,27–28). Targeting the viral protease has been shown as a successful therapeutic strategy for ZIKV as well as the infections caused by other members in the flaviviridae family including Dengue, West Nile, and Hepatitis C viruses (29–34).

**Fig. 1.**
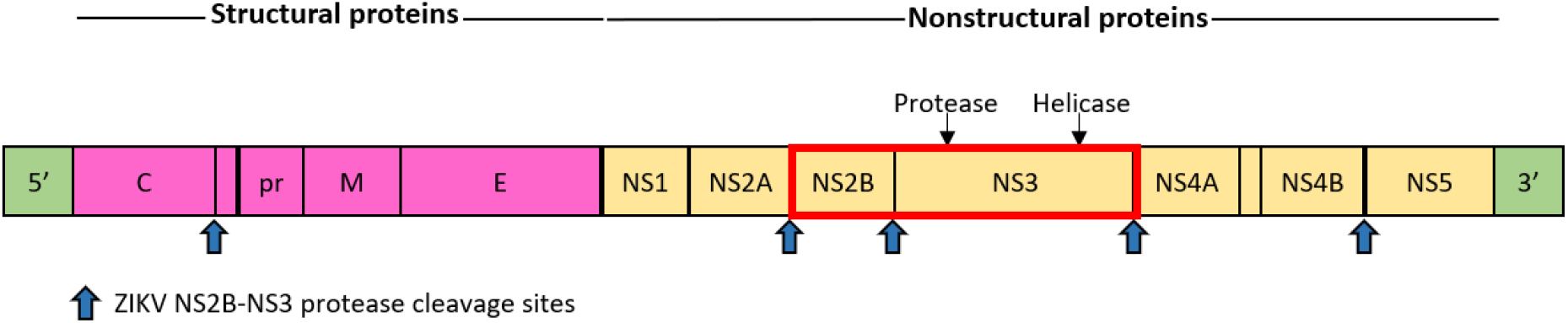
Schematic representation of ZIKV genome. ZIKV polyprotein precursor includes structural proteins, shown in magenta, and nonstructural proteins, shown in yellow. ZIKV NS2B-NS3pro, highlighted by a red box, has an important role for viral protein activation and its cleavage sites are represented by blue arrows.

Designing or screening for effective competitive inhibitors of flavivirus proteases has been challenging since strong and specific binding to S1 and S2 active sub-pockets requires hydrophilic and electrostatic interactions, usually associated with low membrane permeabilities (30). Therefore, we sought to identify a novel type of noncompetitive inhibitors of NS2B-NS3pro that prevent formation of active ZIKV NS2B-NS3pro complex. We chose to target a recently identified transient pocket in the so called “super-open” conformation, described in (35). Here we employed a large scale *in-silico* docking screen, utilizing the Molsoft-ICM software, of approximately 7 million compounds to find effective non-covalent inhibitors for the allosteric pocket of ZIKV NS2B-NS3pro. Seven selected top candidates were tested experimentally, and six of them were confirmed to inhibit enzymatic activity of the viral protease.

## MATERIALS AND METHODS

### Reagents

Routine laboratory reagents were purchased from Thermo Fisher Scientific. The tested compounds were purchased from Chembridge Corp.

### Cloning and purification of ZIKV NS2B-NS3pro construct

A recombinant construct expressing ZIKV NS2B-NS3 protease with 6xHisTag on its N-terminus was used to transform competent *E. coli* BL21 (DE3) Codon Plus cells obtained from Stratagene. Transformed cells were grown at 30°C in LB broth containing carbenicillin (0.1 mg/ml). Cultures were induced with 0.6 mM IPTG for 16 h at 18°C. Cells were collected by centrifugation, re-suspended in Tris-HCl buffer, pH 8.0, containing 1 M NaCl, and disrupted by sonication (30 sec pulse, 30 sec intervals; 8 pulses) on ice. The pellet was removed by centrifugation (40,000xg; 30 min). The construct was then purified from the supernatant fraction on a Ni-NTA Sepharose, equilibrated with 20 mM Tris-HCl buffer, pH 8.0, supplemented with 1 M NaCl. After washing out the impurities using the same buffer supplemented with 35 mM imidazole, the bound material was eluted using a 35 mM - 500 mM gradient of imidazole. The fractions containing recombinant protein were combined, dialyzed against 20 mM Tris-HCl, pH 8.0, containing 150 mM NaCl. The purified material was kept at - 80°C until use. The purity of the material was tested by SDS gel-electrophoresis (12% NuPAGE-MOPS, Invitrogen) followed by Coomassie staining and by Western blotting with anti-HisTag antibodies.

### Proteinase activity assay with fluorescent peptide

The peptide cleavage activity assay with the purified ZIKV NS2B-NS3pro samples was performed in 0.2 ml 20 mM Tris-HCl buffer, pH 8.0, containing 20% glycerol and 0.005% Brij-35. The cleavage peptide pyroglutamic acid Pyr-Arg-Thr-Lys-Arg-7-amino-4-methylcoumarin (Pyr-RTKR-AMC) and enzyme concentrations were 20 μM and 10 nM, respectively. Reaction velocity was monitored continuously at λex = 360 nm and λex = 465 nm on a Tecan fluorescence spectrophotometer (**Tecan Group Ltd.,** Männedorf, Switzerland). All assays were performed in triplicate in wells of a 96 well plate.

### Determination of the IC_50_ values of the inhibitory compounds

The ZIKV NS2B-NS3pro construct (20 nM) was pre-incubated for 30 min at 20°C with increasing concentrations of the individual compounds in 0.1 ml 20 mM Tris-HCl buffer, pH 8.0, containing 20% glycerol and 0.005% Brij 35. The Pyr-RTKR-AMC substrate (20 μM) was then added in 0.1 ml of the same buffer. All assays were performed in triplicate in wells of a 96-well plate. IC_50_ values were calculated by determining the concentrations of the compounds needed to inhibit 50% of the NS2B-NS3pro activity against Pyr-RTKR-AMC. Fitting was performed using GraphPad Prism software.

### Virtual Ligand Screening of the compound library

The structure of NS2-NS3pro was obtained from X-ray crystal structure (PDB ID 5TFN) (35). The “super-open” conformation and the transient pocket exposed in that conformation were identified by comparison of 5LC0 (closed) with five other structures (PDB ID: 5TFN, 5TFO, 6UM3, 5T1V, and 2GGV) (24, 35–37). The single chain construct in 5TFN contained both NS2 and NS3 domains, and the NS2 domain was marked to map possible interactions of selected inhibitors with the NS3 chain. A docking screen was performed against approximately 7 million small molecules from the eMolecules catalog (38). The scored binding poses of small molecules were predicted by the Biased-Probability Monte Carlo optimizer in Molsoft ICM software, and binding affinity of small molecules to the receptor were ranked based on force field based docking score extended with additional free-energy contributions. All scoring functions and pharmacokinetic properties prediction were performed using Molsoft ICM-Pro v3.9 (39–41).

## RESULTS AND DISCUSSION

ZIKV NS2B-NS3 protease is a two-component chymotrypsin-like serine protease consisting of the NS2B co-factor and an NS3 protease domain, similar to other *Flaviviridae* members. The active site of ZIKV NS2B-NS3pro includes three conserved amino acid residues forming a classic catalytic triad (His51, Asp75, and Ser135 in ZIKV) (17). Because of its importance to the virus life cycle and propagation, NS2B-NS3pro is a promising target for antiviral drug design. Unfortunately, due to its shallow, solvent-exposed pocket, as well as the high structural homology of its active center to those of multiple cellular serine proteases, the development of inhibitors with high cellular permeability, stability, and selectivity which target at the active site is challenging (42). As such, the design of inhibitors targeting new allosteric sites on ZIKV NS2B-NS3pro presents an attractive option to successfully avoid this obstacle and develop effective drugs and therapeutic approaches.

### Identification of allosteric site on ZIKV NS2B-NS3 protease

Recently, four structures of ZIKV NS2B-NS3 protease were deposited to the protein data bank (PDB 5TFN, 5TFO, 6UM3 and 5T1V) (24). The protease in all four structures contained a novel “super-open” pocket configuration. In these structures the “closed” conformation observed in PDB ID 5LC0 (36) undergoes a transition to the super-open conformation and reveals a targetable pocket. This transient pocket may be present in the proteases of other flaviviruses (e.g. PDB ID 2GGV for the West Nile virus protease) (35). These novel structures demonstrate the detachment of the C-terminal part of NS2B from a deep pocket. *Our hypothesis was that the major reorganization of the NS3pro C-terminal loop creates a transient novel druggable pocket that can be used for the development of specific binding disrupting the protease activity allosterically* (**Fig. 2A**). The list of contact residues, W83, L85, V146, A87, A88, I147, G148, D86, L149, V154, V155, I123, D120, T118, G153, N152, Q74, D75 and, L76, in the targeted pocket has favorable characteristics for drug-like small molecules, such as shape, depth and hydrophobicity. All structural rearrangements of ZIKV protease in the “super-open” conformation are incompatible with the catalytic activity of the protease. However, this catalytic activity of NS2B-NS3pro can be completely restored via transitioning back to the “closed” conformation. It was unknown experimentally if this pocket could be targeted by a small molecule because of its transient nature, but the pocket's attractive characteristics motivated us to find small molecule binders by a structure-based docking screen.

**Fig. 2.**
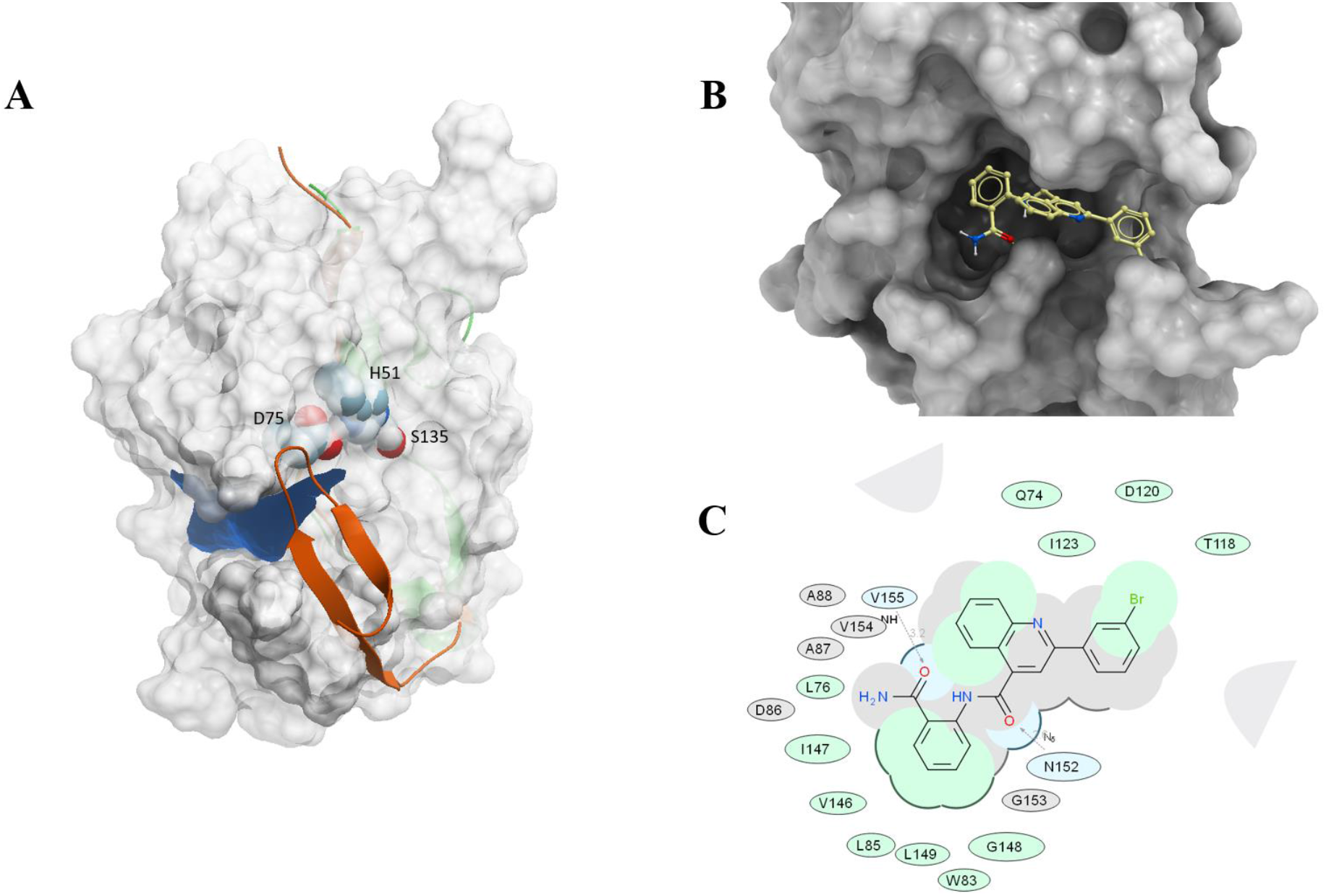
Targeting a hidden pocket exposed in the “super-open” conformation of ZIKV NS2B-NS3 protease. **A.** ZIKV NS3 subunit in grey, active site in purple, beta hairpin of NS2B shown in closed conformation, opens a transient (allosteric) pocket by moving away into an open (green ribbon) conformation. Targeted allosteric pocket is in blue. Catalytic triad of NS3, S135, H51, D75, are labeled and shown in CPK; **B.** Predicted 3D binding pose of RI07 targeting the super-open conformation of NS2B-NS3 protease; **C.** 2D interaction diagram for predicted pose of RI07 compound (W83, L85, V146, A87, A88, I147, G148, D86, L149, V154, V155, I123, D120, T118, G153, N152, Q74, D75, L76). Hydrogen bonding interactions are shown by grey dotted lines.

### Virtual Docking Screen of seven million compounds targeting the allosteric hidden site of ZIKV NS2B-NS3 protease

The structural model of the “super-open” state of the ZIKV protease (PDB 5TFN) was docked and scored against approximately 7 million small molecules from the eMolecules database. Chemicals were first filtered by removing compounds with a high toxicity propensity based on over 1000 structural alerts (41). The binding free energy of each compound to the allosteric pocket was estimated by a docking score computed via the MolSoft ICM-Pro package (39–40). Ten different classes of compounds with the top ICM-Docking scores were initially tested in ZIKV NS2B-NS3 protease enzymatic assay described in the Methods section.

### Selection of core functional group

Ten protease inhibitor candidates suggested from predicted docking poses and binding scores were purchased from available vendors and tested against 10 nM of ZIKV NS2B-NS3 in proteolytic activity assays with 20 μM of fluorogenic peptide substrate, Pyr-RTKR-AMC, in order to validate the viral protease inhibition. We found that the compounds with phenylquinoline and aminobenzamide groups demonstrated desirable activity indicating a potential scaffold for further optimization of allosteric inhibitors of ZIKV NS2B-NS3 protease.

### Variation of phenylquinoline and aminobenzamide substituents and enzymatic activities against ZIKV NS2B-NS3 protease

Phenylquinoline and aminobenzamide-containing compounds showed high affinity to ZIKV NS2B-NS3 in proteolytic activity assays. That prompted us to search for more potent inhibitors from the same chemical class. The binding conformation of RI07, used as a representative phenylquinoline and aminobenzamide substituent compound, in the deep allosteric pocket of super-open conformation of NS2B-NS3 is shown in **Fig. 2B** and **C** suggesting the strong binding affinity between aminobenzamide and hydrophobic residues including W83, L85, D86, A87, A88, V146, and L149 in the identified allosteric pocket of ZIKV NS2B-NS3. An extended set of compounds of various substituents on phenylquinoline was identified in a chemical vendor catalog to test in the same inhibition assay. This iteration resulted in several improved inhibitors labelled RI07, RI22, RI23, RI24, RI27, and RI28.

In addition, the relevant pharmacokinetic and toxicity properties, including cells permeability, water solubility, propensity for Pan-assay interference, potassium channel blocking activity, and toxicity of all candidates was evaluated using appropriate Molscreen models from MolSoft ICM-Pro package (39–41). The predicted parameters were in permissible ranges and are shown in **Table S1**. The docking poses of all six compounds in this series, found in **Fig. 1S**., showed consistent binding conformations in the identified allosteric pocket. The novel compounds in this series were tested in the proteolytic activity assay and stronger inhibitors of ZIKV protease were identified (**Fig. 3.**) with the **IC50** ranging in low micromolar from 3.8 to 14.4 μM. These results show the possibility of targeting the allosteric pocket which is necessary for ZIKV NS2B-NS3 complex formation and the high potential of the compounds containing phenylquinoline and aminobenzamide groups for further development as anti-flavivirals.

**Fig. 3.**
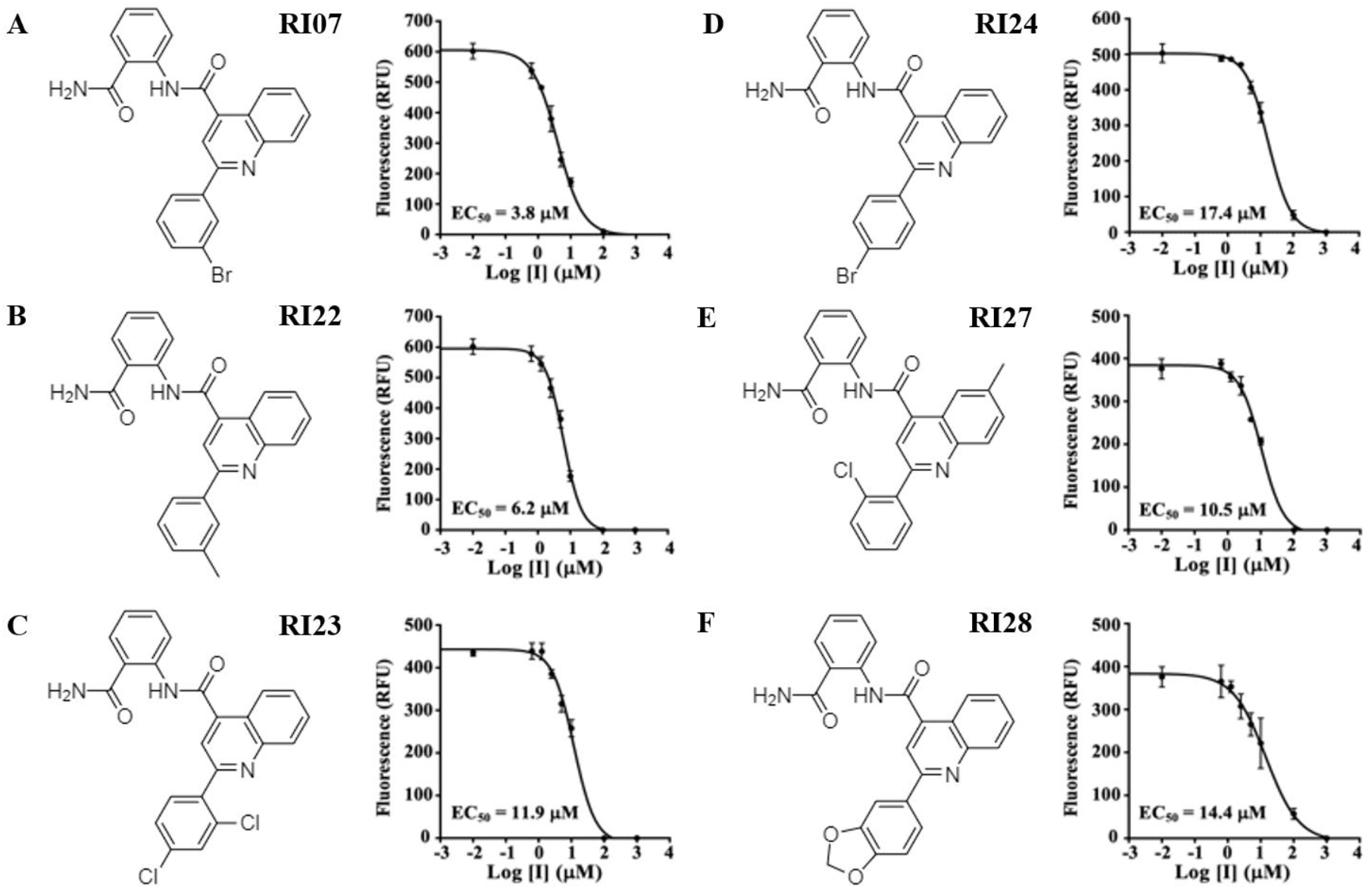
Six experimentally validated inhibitors of catalytic activity of Zika virus protease and their dose response curves. The dose-response curves showing inhibition of NS2B-NS3 protease by RI07, RI22, RI23, RI24, RI27, and RI28 are shown next to the chemical structures. The EC_50_ values of the compounds against ZIKV NS2B-NS3 protease were measured in a protease cleavage assay with Pyr-Arg-Thr-Lys-Arg-MCA fluorescent peptidic substrate (Peptides International - MRP-3159-v). GraphPad Prism was used for curve-fitting. A-F. The X axis shows decimal logarithms of compound concentrations in micromolar units vs. the peptidic substrate cleavage in PFUs.

### Antiviral activity in cells-based assay and cell viability evaluations

The antiviral activities of six identified ZIKV NS2B-NS3 allosteric inhibitors were evaluated using immunofluorescence signal to visualize the number of ZIKV copies in human Neural progenitor cells (NPCs). The antiviral activities of each compound at 10 μM were measured as a ratio between fluorescence signals from the ZIKV envelope protein and cell nuclei represented by DAPI. At 10 μM, all six candidates exhibited significant viral reduction, from approximately 10 to 100-fold compared to 0.1% DMSO chosen as the negative control (**Fig. 4A.**). Additionally, the cell viability of the NPCs at the presence of 10 μM of all six compounds was measured as DAPI signal intensities to evaluate potential cytotoxic activity (**Fig 4B**). All six compounds displayed excellent antiviral activities and low toxicity at 10 μM. Compounds RI22 and RI27 showed slightly higher antiviral activities compared to other four hits due to the methyl substitution on phenylquinoline and aminobenzamide, respectively. Structural analysis shows that these modifications could help enhance hydrophobic interaction with the pocket. These findings confirmed our hypothesis that this novel allosteric pocket, revealed transiently in “super-open” conformation, can be used as a target for a new type of antivirals for Zika and other *Flaviviridae*. This target is beneficial because of its conservation, and hence its lower propensity for treatment escape mutations, and its favorable shape. The identified allosteric inhibitors targeting ZIKV NS2B-NS3 protease and preventing viral proliferation may be further optimized for improved efficacy, PK/PD, and reduced adverse side effect profile.

**Fig. 4.**
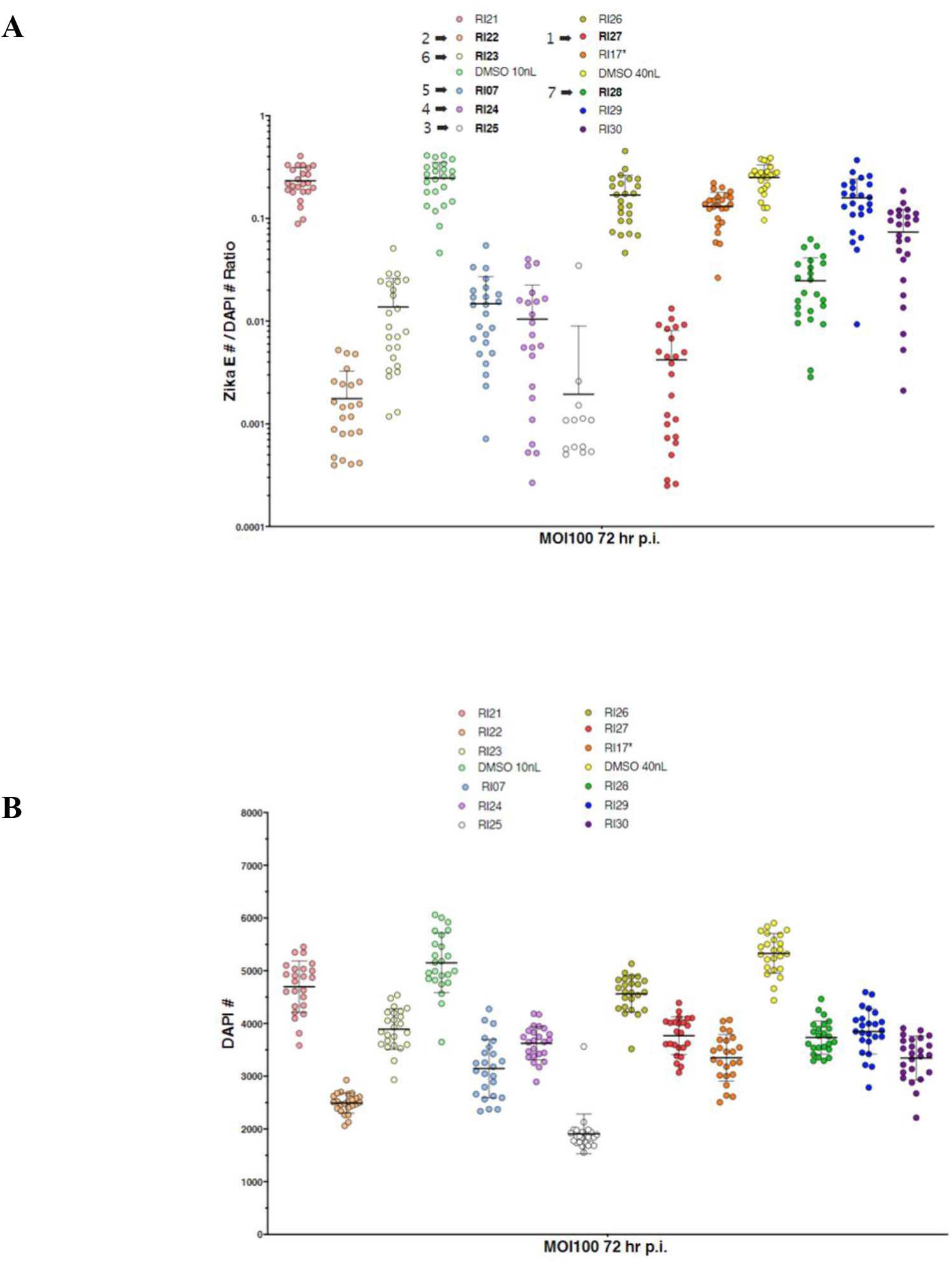
Antiviral activities in ZIKV infected human NPCs cells and cytotoxicity of selected compounds. **A.** The antiviral activities are represented by the immunofluorescence ratio from ZIKV envelope protein and DAPI with the presence of RI07, RI22, RI23, RI24, RI27, and RI28 at 10 μM compared to 0.1% of DMSO. **B.** Cellular cytotoxicity effects were represented by the signal of DAPI with the presence of RI07, RI22, RI23, RI24, RI27, and RI28 at 10 μM.

## CONCLUSIONS

Infections with ZIKV are a current global health concern due to the risk of microcephaly in newborns and Guillain-Barré syndrome in adults. Since there is no approved anti-ZIKV agent and vaccination strategies are limited due to antibody dependent enhancement (ADE) effects observed in Dengue and other flaviviruses. Discovery of small molecule orally available drugs as ZIKV infection treatments has previously been limited. Here we report a new strategy of targeting a super-open conformation of ZIKV NS2B-NS3 protease suggested by analysis of the existing X-ray crystal structures of all flaviviral proteases. The newly identified pocket is a preferable target compared to an active site due its shape, conservation, and hydrophobicity. We have identified six novel compounds containing phenylquinoline and aminobenzamide that inhibited ZIKV NS2B-NS3 protease at IC50 ranging from 3.8 to 14.4 μM. These six compounds were evaluated and validated in cell-based infectivity and toxicity assays. The predicted pharmacokinetic properties of all compounds were also evaluated and proved to be both promising and drug-like. Targeting the identified transient pocket may prove to be a promising strategy for fighting other flaviviral infections as well due to the similar pattern of inactive-to-active state transitions of the viral NS2B-NS3 system.

## ACKNOWLEDGMENTS

We thank Alexander Aleshin for his help, discussions, and X-ray crystallography work which laid the foundation for our screen. We also thank Conall Sauvey for assistance proofreading this manuscript. This work was supported in part by NIH R35GM131881 to R.A., and by NIH R01 NS105969-01 to A.V.T.

## AUTHOR CONTRIBUTIONS

R.A. and A.V.T designed and supervised the study. I.M., S.A.S., Y.W.L., C.H.H. and C.T.H. conducted the experiments. S.A.S., I.M., A.V.T. and R.A. analyzed the data and wrote the manuscript.

## COMPETING FINANCIAL INTERESTS

The authors declare no competing financial interests.

## Notes

### Competing Interest Statement

The authors have declared no competing interest.

