## Supplemental Table 1 and Supplemental Figure 1. for "Allosteric inhibitors of Zika virus NS2B-NS3 protease targeting protease in super-open conformation": Allosteric inhibitors of Zika virus NS2B-NS3 protease_supplementary data.pdf

**Table S1.** Pharmacokinetic parameters, and binding score against ZIKV NS2B-NS3 protease for all compounds.

| Compounds ID | Biding Score[a] | Predicted Pharmacological Properties[b] | | | | | | |
|---|---|---|---|---|---|---|---|---|
| | | molLogS[c] | drugLikeness[d] | molCACO2[e] | molPAMPA[f] | molHERG[g] | molPAINS[h] | Tox_Score[i] |
| RI07 | -37 | -5.12 | 0.57 | -5.09 | -4.71 | 0.13 | 0.05 | 0 |
| RI22 | -31 | -4.50 | 0.59 | -4.99 | -4.71 | 0.11 | 0.01 | 0 |
| RI23 | -31 | -5.92 | 0.70 | -5.08 | -4.80 | 0.37 | 0.06 | 0.42 |
| RI24 | -30 | -5.54 | 0.93 | -5.07 | -4.70 | 0.09 | 0.04 | 0 |

| | | | | | | | | |
|---|---|---|---|---|---|---|---|---|
| RI27 | -33 | -4.65 | 0.78 | -5.10 | -4.63 | 0.09 | 0.04 | 0.42 |
| RI28 | -29 | -4.45 | 0.40 | -5.26 | -4.61 | 0.07 | 0.02 | 0 |

[a]Binding score was calculated using Dockscan function in ICM-Pro (23).

[b]The prediction of relevant pharmacological properties were calculated using Chemical Properties prediction function in ICM-Pro (41).

[c]Water solubility (molLogs) in 10-based logarithm of the solubility in M.

[c]Druglikeness, value range between -1 to 1 higher number indicates more drug-like properties.

[d]CACO-2 permeability, value over -5 indicates high permeability.

[f]PAMPA permeability, value over -5 indicates high permeability.

[g]hERG inhibition, value over 0.5 indicates high probability of being hERG inhibitor.

[h]Pan Assay Interference Compound (PAINS), value over 0.5 indicates high probability of being PAIN compound.

[i]Chemical Alert collected from chemical supplier and other sources, value over 1 indicates molecule contain unfavorable substructure or substituent and being toxic compound.

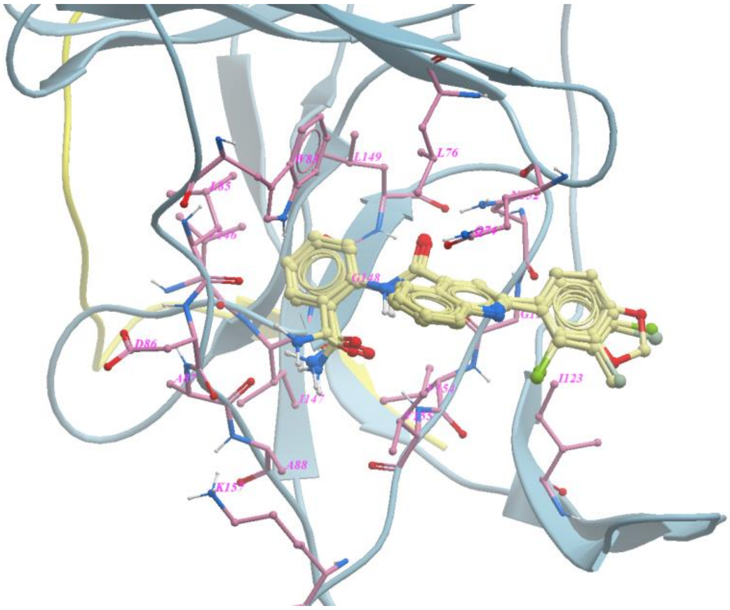


**Figure S1.** Predicted binding conformations of RI07, RI22, RI23, RI24, RI27, and RI28 targeting the open conformation of NS2B-NS3 protease at the identified allosteric pocket.
